# Insular cortex corticotropin-releasing factor integrates stress signaling with social decision making

**DOI:** 10.1101/2021.03.23.436680

**Authors:** Nathaniel S. Rieger, Juan A. Varela, Alexandra Ng, Lauren Granata, Anthony Djerdjaj, Heather C. Brenhouse, John P. Christianson

## Abstract

Impairments in social cognition manifest in a variety of psychiatric disorders, making the neurobiological mechanisms underlying social decision making of particular translational importance. The insular cortex is consistently implicated in stress-related social and anxiety disorders, which are associated with diminished ability to make and use inferences about the emotions of others to guide behavior. We investigated how corticotropin releasing factor (CRF), a neuromodulator evoked by both self and social stressors, influenced the insula. In acute slices from male and female rats, CRF depolarized insular pyramidal neurons. In males, but not females, CRF suppressed presynaptic GABAergic inhibition leading to greater excitatory synaptic efficacy in a CRF receptor 1 (CRF_1_) and cannabinoid receptor 1 (CB_1_) dependent fashion. In males only, insular CRF increased social investigation, and CRF_1_ and CB_1_ antagonists interfered with social decision making. To investigate the molecular and cellular basis for the effect of CRF we examined insular CRF_1_ and CB_1_ mRNAs and found greater total insula CRF_1_ mRNA in females but greater CRF_1_ and CB_1_ mRNA colocalization in male insular cortex glutamatergic neurons which suggest complex, sex-specific organization of CRF and endocannabinoid systems. Together these results reveal a new sex-specific mechanism by which stress and affect contribute to social decision making.

## INTRODUCTION

Stressors and other salient emotional stimuli trigger a shift in attention and cognitive resources that orient decision making and organize situationally adaptive behaviors. In the brain, the structure that mediates the transition between resting and executive cognitive networks is the insular cortex ^1,2^. The insula is anatomically situated to integrate sensory with emotional and cognitive processes ^3,4^. Not surprisingly, insula is associated with many cognitive functions and, in human neuroimaging studies, insula activity correlates with emotion recognition, pain, drug craving and anticipatory fear ^5–8^. Aberrant activity and functional connectivity of the insula and associated network structures leads to hypervigilance, increased interoception and poor emotion regulation, hallmark symptoms of many neuropsychiatric disorders including autism spectrum disorders, schizophrenia, and posttraumatic stress disorder ^9^.

Stress is a major precipitating factor for mental illness. Both self and social stressor exposure initiate the hypothalamic-pituitary-adrenocortical axis response by activation of corticotropin-releasing factor (CRF) neurons in the paraventricular hypothalamus (PVN) ^10^. The CRF system is complex, consisting of 2 receptor subtypes (CRF_1_ and CRF_2_) which couple to a variety of G-proteins expressed in brain region, cell type, and sex specific ways ^11,12^. CRF_1_ receptors and CRF immunoreactive fibers are expressed throughout the corticolimbic system, including in the insular cortex ^13–16,16^. Observing others in distress is highly salient and a potent driver of insular activity which is thought to contribute to empathic cognition ^17,18^. A fundamental precursor to empathy is emotion contagion, a primitive process by which the affective state of a demonstrator leads to a complementary state in the observer ^19^. CRF neurons of the PVN are activated upon exposure to stressed conspecifics which is a mechanism providing for the social transfer of stress responses ^10^. CRF may shape social behaviors by actions at CRF receptors located among the distributed network of neural structures, including the insular cortex, that are engaged by social stress signals and organize social behavior ^20^.

Seeing CRF as a putative modulator of insula cortex functioning and considering the significant influence of stress on psychosocial processes, we investigated the effects of CRF on insular cortex physiology and social behavior. In whole-cell recordings of insular cortex pyramidal neurons, CRF depolarized the membrane potential. This translated to an increase of excitatory synaptic transmission, but only in recordings from male rats. The gain of synaptic efficacy appeared to be a case of CRF causing depolarization induced suppression of inhibition^21^ (DSI) as the effects of CRF were dependent on both GABA_A_ and cannabinoid type 1 receptor (CB_1_). In social behavior, CRF augmented social investigation while a CRF_1_ antagonist interfered with social interactions with stressed conspecifics in male but not female rats. Because we observed sex specific effects of CRF on physiology and behavior, we hypothesized that sex differences exist in CRF_1_ and CB_1_ at the cellular and molecular levels. We employed a combination of quantitative (qPCR) and anatomical (*in situ* hybridization) analyses and found sex differences in CRF_1_ mRNA expression and differences in the cellular distribution of the transcripts in the insular cortex. Together, the data lead us to conclude that CRF, acting upon CRF_1_ receptors depolarizes pyramidal neurons triggering the release of endocannabinoids which suppress presynaptic inhibition. The result is facilitated flow of information through the insula which appears to be necessary for coordinating social interactions with stressed conspecifics.

## RESULTS

### Corticotropin-releasing factor depolarizes insular cortex pyramidal neurons

To determine the effects of CRF on insular pyramidal neurons we first used whole cell current-clamp recordings in acute slices from male and regularly cycling female adult rats. Active and passive intrinsic properties (Table 1) were computed from 14 male and 12 female neurons before and after application of CRF (50nM). Each parameter was analyzed for sex differences, CRF effects and sex by CRF interactions. With regard to sex differences in intrinsic properties, male and female intrinsic properties under aCSF recording conditions were comparable; no main effects of stress were present for any parameter. CRF altered many intrinsic properties including depolarization of resting membrane potential (Fig. 1A), reduction in action potential (AP) amplitude and rise rate (Fig. 1B), and corresponding increase in AP half-width (Fig. 1C). Many insular cortex pyramidal neurons have a bursting phenotype ^22^ and CRF appeared to reduce after depolarization (ADP) amplitude (Fig. 1D) and increased the current required to elicit a burst (Burst Ratio, Fig. 1E). Turning to passive properties, a few sex-specific effects of CRF emerged. In male neurons, CRF reduced input resistance (Fig. 1F) and in female neurons, CRF reduced the membrane time constant. Interestingly, CRF increased rectification ratio (Fig. 1G) to a similar extent in males and females.

**Table 1.** Effect of 50nM CRF on intrinsic properties of insular cortex pyramidal neurons.

| Parameter | Sex | Means (SEM) |  | Results |  | Summary |
| --- | --- | --- | --- | --- | --- | --- |
|  |  | aCSF | CRF | Main effects | Interaction |  |
| Membrane potential (mV) | M | -66.65<br>(1.527) | -59.64<br>(2.007) | CRF, $F_{1,24} = 72.93$, $p < 0.0001$ | $F_{1,24} = 5.124$,<br>$p = 0.033$ | ↑ |
| | F | -68.41<br>(1.189) | -64.33<br>(1.268) | Sex, $F_{1,24} = 2.267$,<br>$p = 0.145$ | | ↑ |
| Rheobase (pA) | M | 1256.42(<br>105.92) | 1226.42<br>(129.94) | CRF, $F_{1,24} = 0.282$,<br>$p = 0.600$ | $F_{1,24} = 0.924$,<br>$p = 0.346$ | = |
| | F | 1515.00(<br>132.76) | 1619.17(<br>157.49) | Sex, $F_{1,24} = 3.573$,<br>$P = 0.070$ | | = |
| Spike Amplitude (mV) | M | 73.35<br>(3.852) | 70.99<br>(4.039) | CRF, $F_{1,24} = 4.933$, $p = 0.036$ | $F_{1,24} = 0.263$,<br>$p = 0.612$ | ↓ |
| | F | 80.58<br>(3.764) | 76.80<br>(4.136) | Sex, $F_{1,24} = 1.430$,<br>$p = 0.243$ | | ↓ |
| AP Rise Rate (mV/ms) | M | 181.1<br>(17.55) | 157.6<br>(14.548) | CRF, $F_{1,24} = 18.93$, $p = 0.0002$ | $F_{1,24} = 0.535$,<br>$p = 0.472$ | ↓ |
| | F | 229.2<br>(25.536) | 196.2<br>(26.136) | Sex, $F_{1,24} = 2.253$,<br>$p = 0.146$ | | ↓ |
| AP Half-Width (ms) | M | 1.248<br>(0.075) | 1.357<br>(0.064) | CRF, $F_{1,24} = 16.69$, $p = 0.0004$ | $F_{1,24} = 0.134$,<br>$p = 0.717$ | ↓ |
| | F | 1.125<br>(0.074) | 1.256<br>(0.098) | Sex, $F_{1,24} = 1.121$,<br>$p = 0.300$ | | ↓ |
| Afterdepolarization (mV) | M | 10.36<br>(1.365) | 8.593<br>(1.312) | CRF, $F_{1,24} = 26.83$, $p < 0.0001$ | $F_{1,24} = 2.142$,<br>$p = 0.156$ | ↓ |
| | F | 12.98<br>(0.805) | 9.808<br>(0.688) | Sex, $F_{1,24} = 1.554$, $p = 0.225$ | | ↓ |
| Burst Ratio | M | 0.2608<br>(0.052) | 0.3889<br>(0.055) | CRF, $F_{1,21} = 38.82$, $p < 0.0001$ | $F_{1,21} = 3.769$,<br>$p = 0.066$ | ↑ |
| | F | 0.2153<br>(0.032) | 0.2825<br>(0.024) | Sex, $F_{1,21} = 1.600$,<br>$p = 0.220$ | | ↑ |
| Input Resistance (MΩ) | M | 130.9<br>(15.318) | 120.9<br>(14.663) | CRF, $F_{1,24} = 5.985$,<br>$p = 0.0221$ | $F_{1,24} = 2.497$,<br>$p = 0.127$ | ↓ |
| | F | 104.9<br>(13.460) | 102.8<br>(14.249) | Sex, $F_{1,24} = 1.156$,<br>$p = 0.293$ | | = |
| Time Constant (ms) | M | 21.71<br>(0.946) | 20.07<br>(0.946) | CRF, $F_{1,24} = 12.37$, $p = 0.0018$ | $F_{1,24} = 0.173$, $p = 0.681$ | = |
| | F | 24.25<br>(0.799) | 22.17<br>(0.726) | Sex, $F_{1,24} = 4.235$,<br>$p = 0.051$ | | ↓ |
| Rectification Ratio (mV) | M | 0.8343<br>(0.023) | 0.9014<br>(0.023) | CRF, $F_{1,24} = 35.28$,<br>**$p < 0.0001$** | $F_{1,24} = 0.611$<br>$p = 0.442$ | ↑ |
| | F | 0.8158<br>(0.032) | 0.9033<br>(0.025) | Sex, $F_{1,24} = 0.058$,<br>$p = 0.810$ | | ↑ |
| Sag Ratio | M | 1.359<br>(0.175) | 1.384<br>(0.181) | CRF, $F_{1,24} = 1.339$, $p = 0.259$ | $F_{1,24} = 2.072$,<br>$p = 0.163$ | = |
| | F | 1.555<br>(0.244) | 1.331<br>(0.231) | Sex, $F_{1,24} = 0.065$,<br>$p = 0.801$ | | = |
| sAHP (mV) | M | -655.21<br>(85.958) | -765.43<br>(84.764) | CRF, $F_{1,24} = 3.080$, $p = 0.092$ | $F_{1,24} = 0.517$, $p = 0.479$ | = |
| | F | -659.41<br>(49.605) | -705.58<br>(44.492) | Sex, $F_{1,24} = 0.092$,<br>$p = 0.479$ | | = |
| Firing rate<br>(Spikes/s @250pA) | M | 57.214<br>(17.354) | 51.143<br>(19.000) | *CRF, $F_{1,24} = 9.611$,<br>**$p = 0.005$** | *$F_{1,24} = 0.440$,<br>$p = 0.503$" | ↓ |
| | F | 52.092<br>(16.433) | 38.050<br>(14.649) | *Sex, $F_{1,24} = 0.010$,<br>$p = 0.921$ | | = |

Data were analyzed with 2-way analysis of variance. Summary indicates effect of CRF and Sex on parameter measured. Arrows indicate significant differences between aCSF and CRF conditions ( $ps < 0.05$ , Sidak corrected t-test). \*The distribution of firing rates were negatively skewed so the data were converted with a Log transform prior to analysis with ANOVA.

**Fig. 1.**
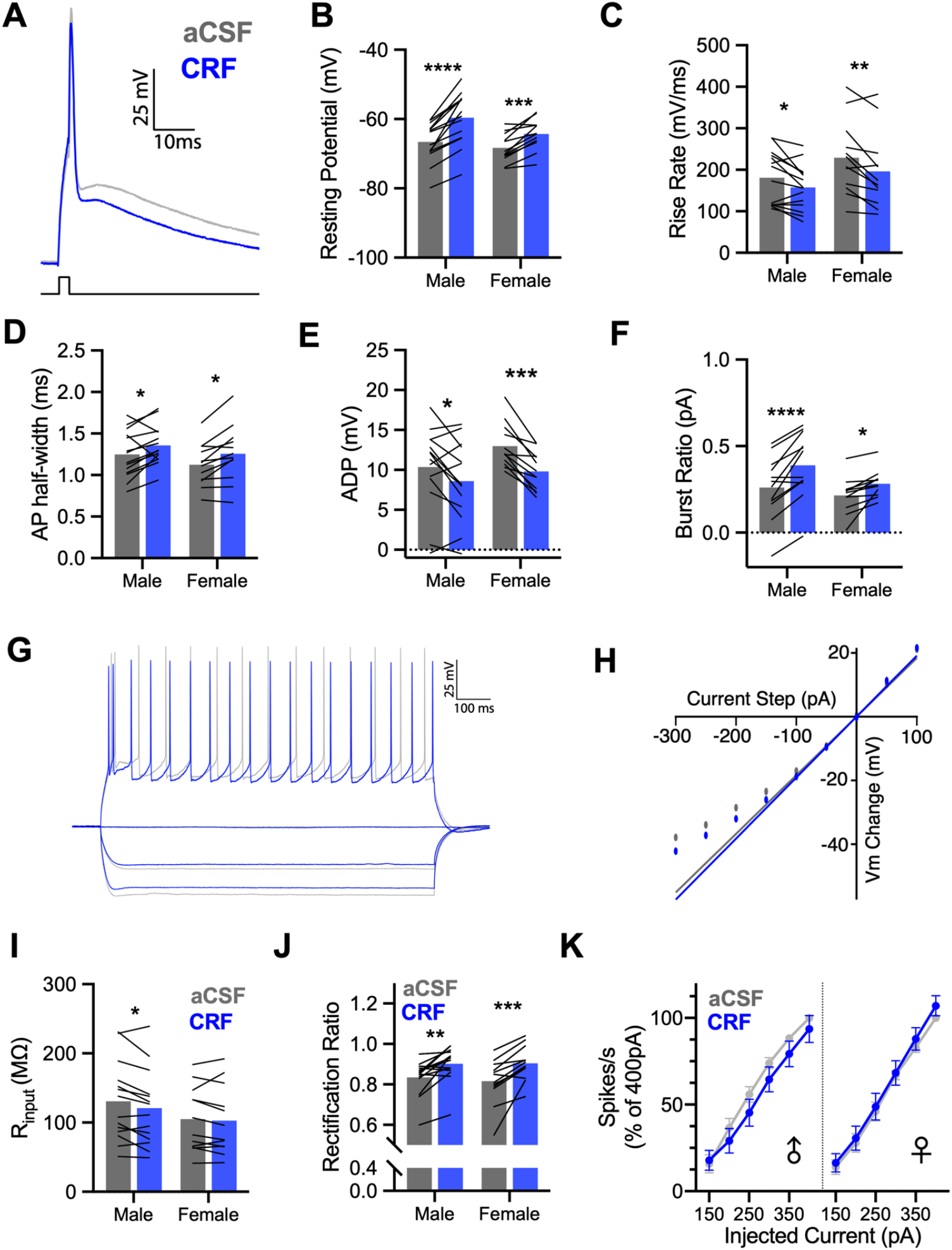
CRF alters intrinsic properties of male and female insular cortex pyramidal neurons in whole cell recordings. **A**. Representative single action potential (AP) recordings of deep layer insular cortex pyramidal neurons at baseline (aCSF-grey) and after application of 50nM Corticotropin-releasing factor (CRF-blue). **B**. CRF decreased the resting potential of male (n = 14) and female (n = 12) pyramidal neurons, F_CRF_(1, 24) = 72.93, P < 0.0001, with this effect being stronger in males than females as indicated by a CRF × sex interaction, F_CRF × SEX_(1, 24) = 5.124, P = 0.033. **C**. Action potential rise rate was reduced by CRF in both males and females, F_CRF_(1, 24) = 18.93, P = 0.0002. **D**. Action potential half-width increased following CRF application in male and female recordings, F_CRF_(1, 24) = 16.69, P = 0.0004. **E**. CRF reduced the amplitude of the after depolarization (ADP) in both male and female recordings, F_CRF_(1, 24) = 26.83, P < 0.0001. **F**. CRF increased the current required to trigger burst firing in male and female neurons, F_CRF_(1, 24) = 38.82, P < 0.0001. **G**. Representative family of 1s hyperpolarizing and depolarizing current injections used characterize passive membrane properties and spike rate in aCSF (grey) and after 50nM CRF (blue). **H**. Example Steady-state current-voltage dependence plot. Input resistance was determined by linear fit and slope at 0pA and deviation from fit indicates rectification. **I**. CRF reduced membrane input resistance in male and female neurons, F_CRF_(1, 24) = 5.985, P = 0.022; this effect appeared most robustly in males. **J**. CRF increased rectification of membrane potential in males and females, F_CRF_(1, 24) = 35.28, P < 0.0001. **K**. CRF did not alter firing rates in response to 1s depolarizing current injections in either males or females. Bar graphs indicate mean with individual replicates, line graphs mean (+/− SEM). *P < 0.05, ** P < 0.01, *** P < 0.001, **** P < 0.0001 (Sidak’s tests).

CRF has high affinity for CRF_1_ and CRF_2_ receptors. Available *in situ* hybridization data and autoradiographic studies of CRF receptor distribution indicated that CRF_1_ is widely expressed in rat cortex, whereas CRF_2_ is only sparsely present or not detected ^13,14^. To establish whether the effects of CRF on insular cortex pyramidal neurons were mediated by action at the CRF_1_ receptor we replicated the intrinsic characterization in 8 adult male neurons. Intrinsic properties were determined after 10 min aCSF, 10 min aCSF with the CRF_1_ antagonist CP154526 (10*μ*M) and then again with CRF (50nM) added. For each parameter that was significantly changed by CRF in the first experiment, the CRF_1_ antagonist appeared to prevent those changes (Sup. Fig 1) suggesting that the CRF_1_ receptor is the primary effector of CRF in the insular cortex.

### CRF augments excitatory neurotransmission in the insula

The effects of CRF on intrinsic properties suggest a mix of augmentation (depolarization) and dampening (reduced AP parameters, reduced bursting) forms of modulation. To better understand how CRF might alter insular cortex information throughput, we investigated CRF effects on synaptic transmission. Using an extracellular multielectrode array (MEA), field excitatory postsynaptic potentials (fEPSP) were characterized in male and female acute slices (Fig. 2A). fEPSP input/output curves were generated in aCSF and then again in either 50 or 300nM CRF from stimulation within the insular cortex. In male, but not female slices, CRF caused a dose-dependent leftward shift indicating augmented synaptic efficacy at both 50 nM (Fig. 2B) and 300 nM (Fig. 3C) concentrations. This is illustrated by an increase in the normalized fEPSP response in males at 300 nM compared to 50 nM which did not occur in female insula slices (Fig. 2D). We then tested whether CRF augmented insular synaptic excitability in males is CRF_1_ dependent by coapplying 300 nM CRF and 10 *μ*M CP154526 during I/O curves. The CRF_1_ antagonist blocked the effects of CRF on fEPSP (Fig. 2F). In summary, CRF application increased excitatory synaptic efficacy in the insula of male rats via action at the CRF_1_ receptor.

**Fig. 2.**
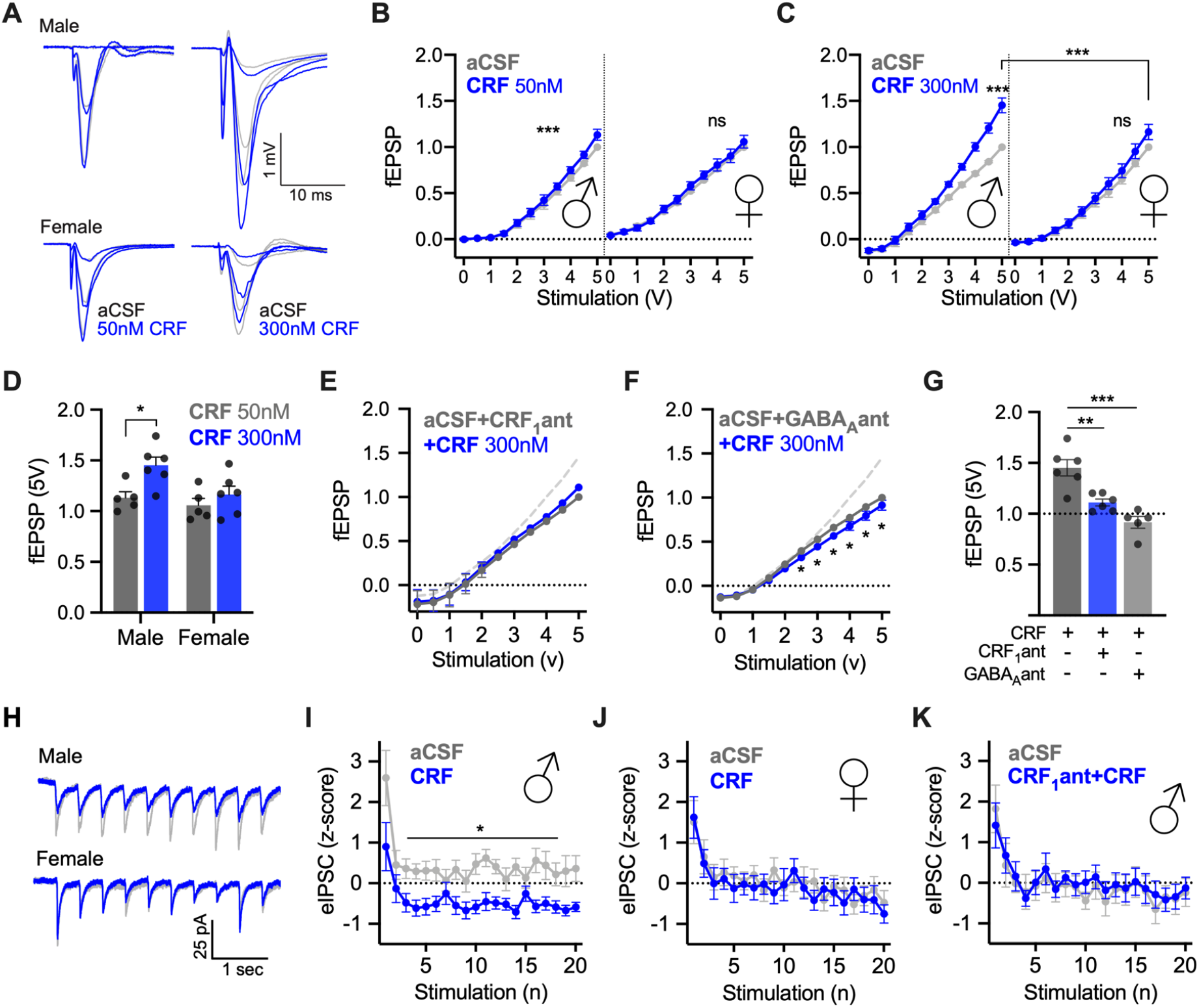
CRF has dose and sex dependent synaptic effects in insular cortex slices. **A**. Representative traces of male (above) and female (below) field excitatory postsynaptic potentials (fEPSP) at 1, 3 and 5 V under aCSF (grey) and after 50 nM (left, blue) or 300 nM CRF (right, blue) conditions. For analysis, traces were normalized to the peak amplitude of the fEPSP evoked at 5V in aCSF. **B**. Bath application of 50nM CRF significantly increased fEPSPs in male insular cortex slices in biphasic 0-5V I/O curves F_Voltage × CRF_ (20, 80) = 5.791, P < 0.0001 with post-hoc tests showing CRF being significantly increased over baseline at 3 V (p = 0.0425), 3.5 V (p < 0.0001), 4 V (p = 0.0001), 4.5 V (P = 0.0091) and 5 V (P = 0.0009). However, there was no significant effect of CRF on female slices F_Voltage × CRF_ (20, 80) = 0.5351, P < 0.5667. A three way ANOVA revealed significant interactions between voltage and CRF and sex: F_Voltage × CRF_(10, 80) =3.654, p = 0.0005, F_Voltage × Sex_(10, 80) = 2.910, p = 0.0037 as well as a main effect of sex, F_Sex_(1, 8) = 10.53, p = 0.0118. **C**. Bath application of 300 nM CRF led to a sex difference in fEPSPs such that males showed increased synaptic efficacy but not females resulting in a significant three way interaction, F_Voltage × Sex × CRF_ (10, 100) = 5.306, p < 0.0001. Males showed significant increases in fEPSP under CRF conditions via Tukey’s multiple comparison tests at 2.5 V (P = 0.0370), 3 V (P = 0.0212), 3.5 V (P = 0.0124), 4 V (P = 0.0063), 4.5 V (0.0040) and 5 V (P = 0.0058). **D**. Comparing 5V responses (normalized to female aCSF 5V) under 50 nM versus 300 nM CRF by sex revealed main effects of Sex, F_Sex_(1, 18) = 5.737, P = 0.0277, and CRF, F_CRF_(1, 18) = 7.855, P = 0.0118. Sidak’s post hoc tests showed that there was a significant dose effect in males t(18) = 2.981, P = 0.0471, but not in females t(18) = 0.9826, P = 0.9165. **E**. CRF_1_ antagonist CP154526 (10 *μ*m) coapplied with 300 nM CRF prevented CRF from increasing fEPSPs in slices from male rats. The dashed grey line depicts the effect of 300nM CRF alone for comparison. While there was a significant interaction F_Voltage × CRF_(20, 89) = 3.276, P < 0.0001. There was no main effect of CRF, F_CRF_(2, 10) = 0.0362, p =0.9646. No post hoc comparisons were significant across treatments at different voltages. **F**. Coapplication of the GABA_A_ antagonist SR95531 prevented the enhancing effect of 300nM CRF and led to a significant decrease in fEPSPs F_Voltage × CRF_(10, 40) = 3.464, P =0.0024 in slices from male rats. Significant Tukey’s post hoc comparisons were found at 2.5V (P = 0.0422), 3V (P = 0.0144), 3. V (p = 0.0038), 4 V (P = 0.0026), 4.5V (P = 0.0014), and 5V (P = 0.0116). **G**. 5V fEPSPs after CRF, CRF+CP154526 and CRF+SR95531 (300nM) were normalized to the relative 5V fEPSP in aCSF in (D) to summarize the effect of CRF_1_ and GABA_A_ receptor antagonist on fEPSP. Both CRF_1_ antagonist and GABA_A_antagonist prevented the increase in fEPSP caused by CRF, F(2, 14) = 19.42, P < 0.0001. Tukey’s post hoc tests show a significant difference between CP154526 + CRF and 300 nM CRF (P = 0.0031) and SR95531 + CRF and 300 nM CRF (P <0.0001). **H**. Voltage-clamp recordings of evoked inhibitory postsynaptic currents (eIPSC) from deep layer insular cortex pyramidal neurons under baseline (grey-aCSF with glutamatergic synaptic antagonists) and after 50nM CRF (blue) in slices from male or female rats. Twenty eIPSCs were evoked by extracellular bipolar electrodes at 5Hz (the first 10 are shown). For analysis, eIPSC amplitudes were normalized using z-scores computed from the mean and standard deviation of the aCSF recordings (Panels I-J). Basal ePISC amplitudes did not differ between male and female recordings. **I**. CRF significantly reduced the amplitude of eIPSCs in males, F_CRF_ (1, 8) = 7.006, P = 0.0294. **J**. CRF did not alter eIPSC amplitudes in females F_CRF_(1, 8) = 0.0531, P = 0.8235. **K**. Pretreatment of the slice with CRF_1_ antagonist eliminated the effect of CRF on eIPSCs in slices from male rats, F_CRF_(1, 7) = 0.0547, P = 0.8218.

**Fig. 3.**
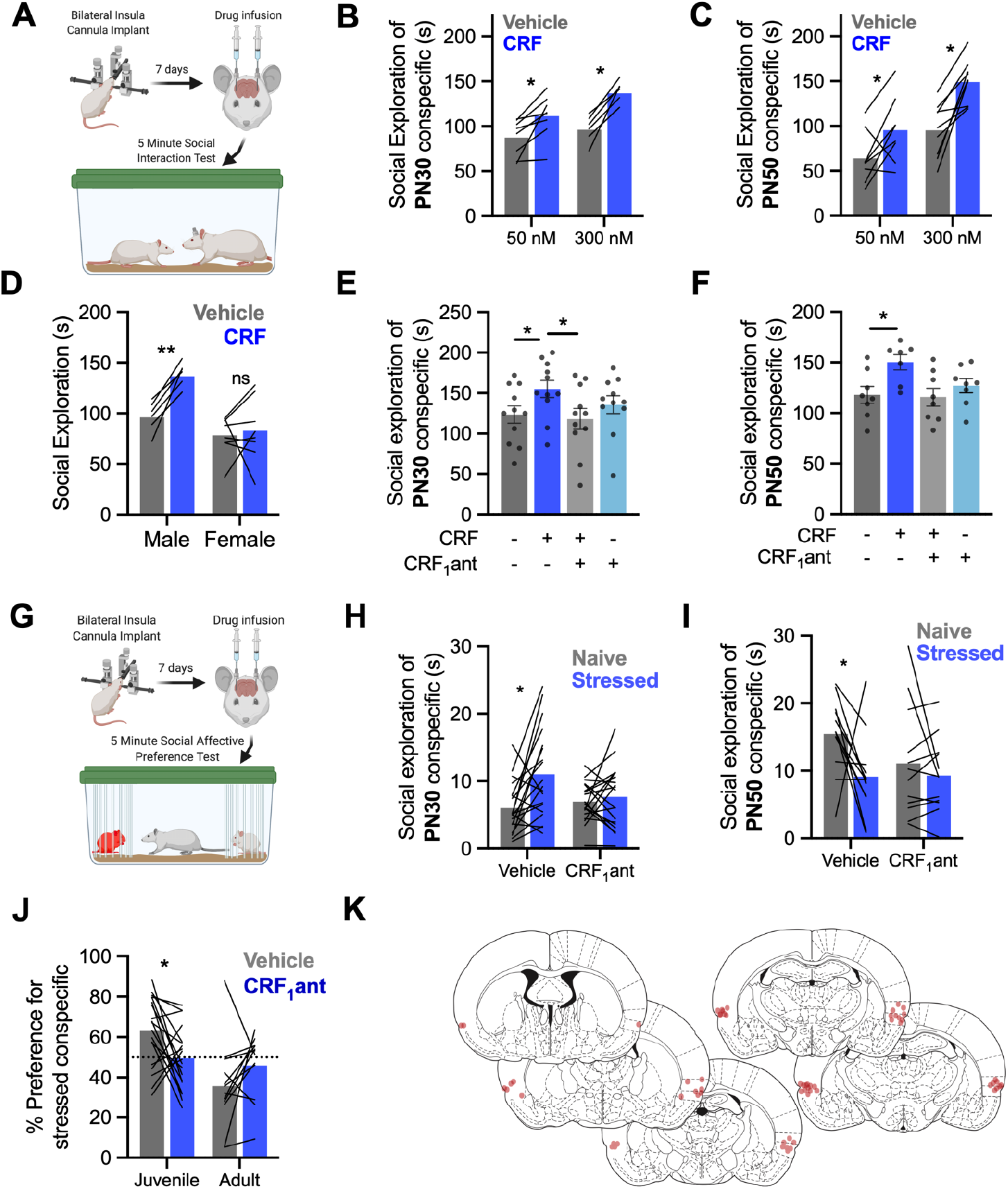
CRF augments social behavior and is necessary for social decision making. **A**. A diagram laying out the experimental procedure for social exploration tests. Cannula were placed in the insular cortex. On the day of testing, rats were given 1h to acclimate to the testing cage. CRF or saline vehicle infusions were made 40 min prior to social interaction with a juvenile (P30) or adult (P50) conspecific for 5 minutes. **B**. In male rats, CRF increased social exploration of juvenile conspecifics, F_CRF_(1, 13) = 48.5, P < 0.0001. Sidak’s post hoc tests revealed significantly increased social exploration at both 50 nM (P = 0.0044) and 300 nM (P < 0.0001). **C**. In male rats, CRF also increased social exploration of P50 conspecifics, F_CRF_(1, 15) = 24.99, P = 0.0002. Sidak’s post hoc tests showed that social exploration was increased at both 50 nM (P =0.0450) and 300 nM (P = 0.0007) doses. **D**. In female rats, 300nM CRF did not alter social interaction with juvenile conspecifics. Female data were compared males at 300nM (data reploted from panel B to facilitate comparison) revealing a sex specific effect of CRF F_Sex × CRF_(1, 13) = 6.517, P = 0.0241 such that males showed increased social exploration following CRF treatment (P = 0.0033) but females did not (P = 0.8615). **E**. Social exploration by males (n = 11) of juvenile (PN30) conspecifics was altered by CRF and the CRF_1_ antagonist CP154526. A 2-way repeated measures ANOVA revealed a significant interaction F_CRF × CRF1antagonist_(1, 10) = 12.82, P = 0.005. 300 nM CRF increased social exploration which was significantly greater than both the vehicle condition (Tukey’s post-hoc test, P = 0.0359) and the combined CRF and CRF_1_ antagonist condition (P = 0.0152). Independently, the CRF_1_ antagonist had no effect on social exploration (P = 0.6093). **F**. In tests of male rats (n = 8) with adult conspecifics, CRF_1_ antagonist blocked the increase in social interaction caused by CRF, F_CRF × CRF1antagonist_(1, 6) = 12.67, P = 0.0119. Mean social interaction time was greatest in the group that received CRF alone which differed from the vehicle (P = 0.0142) and combined CRF and CRF_1_ antagonist conditions (P = 0.0457). **G**. Diagram of the social affective behavior test (SAP) paradigm. Rats received insular cannula implants. On the test day, drug infusions were made 40 min before SAP tests consisting of a 5 minute interaction with a naive and stressed same sex conspecific. The amount of time spent socially investigating each conspecific is recorded. **H**. When tested under vehicle conditions with PN30 conspecifics, male rats n = 19) exhibit greater exploration of the stressed rat (P = 0.0027); this pattern was blocked by the CRF_1_ antagonist (P = 0.8293) supported by a significant interaction, F_CRF1antagonist × Stress_(1, 18) = 5.225, P = 0.0346. **I**. Experimental male rats (n = 11) spent less time interacting with stressed PN50 adult conspecifics in the vehicle condition but this pattern was blocked by the CRF_1_ antagonist, F_CRF1antagonist × Stress_ (1, 10) = 6.133, P = 0.0327. Post hoc comparisons revealed a preference for more interaction with naive adults in vehicles (P = 0.0020) but no difference with the CRF_1_ antagonist (P = 0.5150). **J**. For comparison, time spent interacting with naive and stressed conspecifics from panels H and I was converted to a preference score (% preference = time investigating stressed conspecific / total investigation time * 100). In vehicle conditions, experimental rats preferred interaction with stressed juveniles, but avoided stressed adults and CRF_1_ antagonist treatment appeared to reduce these preferences, F _Age × Drug_ (1, 28) = 11.30, P = 0.0023. When comparing juveniles, the percent preference for the stressed conspecific was significantly reduced by CRF_1_ antagonist (P = 0.0227). When comparing adults, although the CRF_1_ antagonist appeared to eliminate the preference for naive conspecifics, the Sidak corrected posthoc test did not reach significance (P = 0.0768). **K**. Cannula maps showing the placement of in-dwelling cannula across all experiments related to Fig 3. Diagrams in panels A and G were created with *BioRender*.*com*.

### CRF reduces evoked presynaptic GABA release

An enhancement of fEPSP transmission could result from direct effects of CRF on excitatory neurons or from modulation of insular inhibitory tone. Although CRF depolarized intrinsic properties which might predict greater excitatory transmission, it dampened AP size and reduced firing frequency. Therefore, we hypothesized that the enhancement of fEPSP observed after CRF was dependent upon GABA_A_. We tested this by pretreating male insular slices with GABA_A_ antagonist SR95531 (2*μ*M) and then applied CRF (300nM) and repeated fEPSP input/output characterization. In the presence of the GABA_A_ antagonist, CRF had no apparent effect (Fig. 2G). In a direct comparison we found both CRF_1_ antagonist and the GABA_A_ antagonist blocked CRF based increases in insular excitatory synaptic transmission (Fig. 2H) with both antagonists showing similar fEPSPs at 5V (Fig. 3I).

Because it is possible that CRF acts directly on GABAergic neurons in addition to its effects on principle neurons, we next used whole cell, voltage-clamp recordings to investigate the effect of CRF on evoked inhibitory postsynaptic currents (eIPSC) in male and female insular cortex slices. Here, trains of 20 eIPSCs (5Hz) were evoked by a bipolar extracellular electrode placed in close proximity to a pyramidal neuron in a whole-cell recording. The amplitudes of eIPSCs were computed under aCSF and then after application of CRF (50nM). Here, CRF caused a marked reduction in eIPSC amplitude in males (Fig. 2J) but the effect was not present in females (Fig. 2K). Further, pretreatment with the CRF_1_ antagonist eliminated the inhibitory effect of CRF on eIPSCs in males (Fig. 2L). In a separate set of recordings, spontaneous IPSCs were recorded using standard methods and were not sensitive to CRF (Sup. Fig 2).

### Insular CRF increases social exploration of conspecifics in males but not females

The foregoing physiological findings suggest that CRF augments synaptic transmission in IC via downregulation of local GABAergic interneurons. Manipulations that alter insular excitability are associated with changes in social affective behaviors ^22–25^ and the CRF system is implicated in many aspects of social behavior ^26^. To begin to assess whether the effect of CRF in the slices was behaviorally relevant, we utilized social interaction behavior tests. Because fEPSPs were increased in males but not females at 50 and 300 nM doses of CRF we first investigated whether these doses led to functional changes in social investigation towards juvenile (P28) or adult (P50) conspecifics in male rats in a 5 minute social investigation test (Fig. 3A). Bilateral infusion of either 50 nM (238pg/500nL) or 300 nM (1.4ng/500nL) doses of CRF to the insula 40 minutes before social interaction led to an increase in the social investigation of both juvenile (Fig. 3B) and adult (Fig. 3C) conspecifics compared to social investigations following saline injections. To establish whether the sex difference observed in physiology was present in social behavior, a separate cohort of female rats received social interaction tests with juveniles following a 300 nM injection of CRF. Consistent with the physiology, CRF did not alter female social interaction (Fig. 3D).

Next, we tested whether augmentation of male social interaction by CRF was CRF_1_ receptor dependent where adult male rats were given 4 social interaction tests (separate cohorts for juvenile or adult conspecifics) after 0.5 µL/side bilateral infusions of either saline, CRF (300 nM, 1.43ng/µL), CP154526 (10*μ*M, 4.01ng/µL), or a combination of CRF and CP154526. Drug treatment order was counterbalanced and data were initially analyzed for order effects; none were found. As above, CRF increased social interaction compared to saline or CP154526, which was without effect (Fig. 3E-F). Importantly, social interaction levels did not differ from saline control levels when CRF was coadministered with CP154526 showing that CRFs augmentation of social behavior relies on CRF_1_ receptor activation in the insula.

### Social investigation of stressed conspecifics requires insular CRF_1_

The previous gain-of-function results provide evidence that insular CRF and CRF_1_ may contribute to social behavior. In a seminal study, Sterley and colleagues ^10^ demonstrated in mice that the CRF system is robustly activated during social encounters with stressed conspecifics. Similarly, we demonstrated that insula activity determines the nature of social interaction with stressed conspecifics, either approach or avoidance, in a social affective preference (SAP) test ^22,25^. Here, we predicted that upon exposure to a choice to interact with either a stressed or naive conspecific there may be increases in insular cortex CRF release which, via CRF_1_ may contribute to social decision making. The social affective preference (SAP) test exposes test rats to 2 unfamiliar juvenile (PN28) or adult (PN50) conspecifics, one of which is naive and the other which has just received a mild stressor (2, 5-second 1mA footshocks). In the SAP test, adult male rats engage in more social investigation with stressed juveniles, but avoid interaction with stressed adults and this pattern requires the insular cortex.

Male rats were given 5 minute SAP tests with juvenile or adult conspecifics following microinjection of saline or the CRF_1_ antagonist 10*μ*M CP154526 40 minutes prior to testing. CRF_1_ blockade prevented the formation of a preference for stressed juveniles (Fig. 3H) such that approach behavior did not differ towards stressed or unstressed conspecifics. In adult conspecifics (Fig. 3I) control tests indicated a preference for unstressed adult conspecifics but this preference was eliminated via injection of CRF_1_ antagonist. Percent preference for stressed individuals was significantly altered by CRF_1_ injections such that test rats lost preference for both stressed juveniles and non stressed adults (Fig. 3J).

### CB_1_ is necessary for the augmentation of excitatory neurotransmission and social behavior increases caused by CRF

To better resolve the mechanism by which CRF alters insular excitatory/inhibitory tone we considered two possible mechanisms. First, CRF might directly alter GABAergic interneurons, GABA release, or GABA_A_ kinetics. Second, CRF might indirectly modify GABA function via actions that begin with the CRF_1_ receptor on pyramidal neurons. While the former remains interesting, there are technical challenges with direct assessment of interneuron function in rats. Regarding the later, depolarization and accumulation of intracellular calcium leads to the synthesis and retrograde release of endocannabinoids. Via action at the G_i_-protein coupled presynaptic cannabinoid receptor 1 (CB_1_), depolarization leads to reduction in GABA release and the phenomenon of DSI ^21^. Because CRF depolarized pyramidal neurons, we hypothesized that CRF might indirectly affect presynaptic GABA tone via DSI which led us to consider the role of CB_1_ in mediating effects of CRF. We utilized AM251 (2*μ*M), a CB_1_ inverse agonist ^27^, in fEPSPs, eIPSCs and social behavior in male rats. Measuring fEPSPs, we first found that co-application of AM251 prevented CRF effects on synaptic efficacy (Fig. 4A). In whole-cell recordings of eIPSCs, CRF had no effect when administered with AM251 (Fig. 4B). As before, insular CRF injections increased social interaction with juvenile conspecifics. AM251 (Fig. 4C) prevented insular CRF from increasing social investigation of a juvenile conspecific and brought the total amount of social interaction to vehicle levels (Fig. 4D). Turning to SAP tests, AM251 injected into the insula prevented the formation of a preference for stressed juveniles (Fig. 4E) or unstressed adult conspecifics (Fig. 4F) indicating that, in addition to CRF_1_, CB_1_ is necessary for the evaluation of these socioemotional cues (Fig. 4G).

**Fig. 4.**
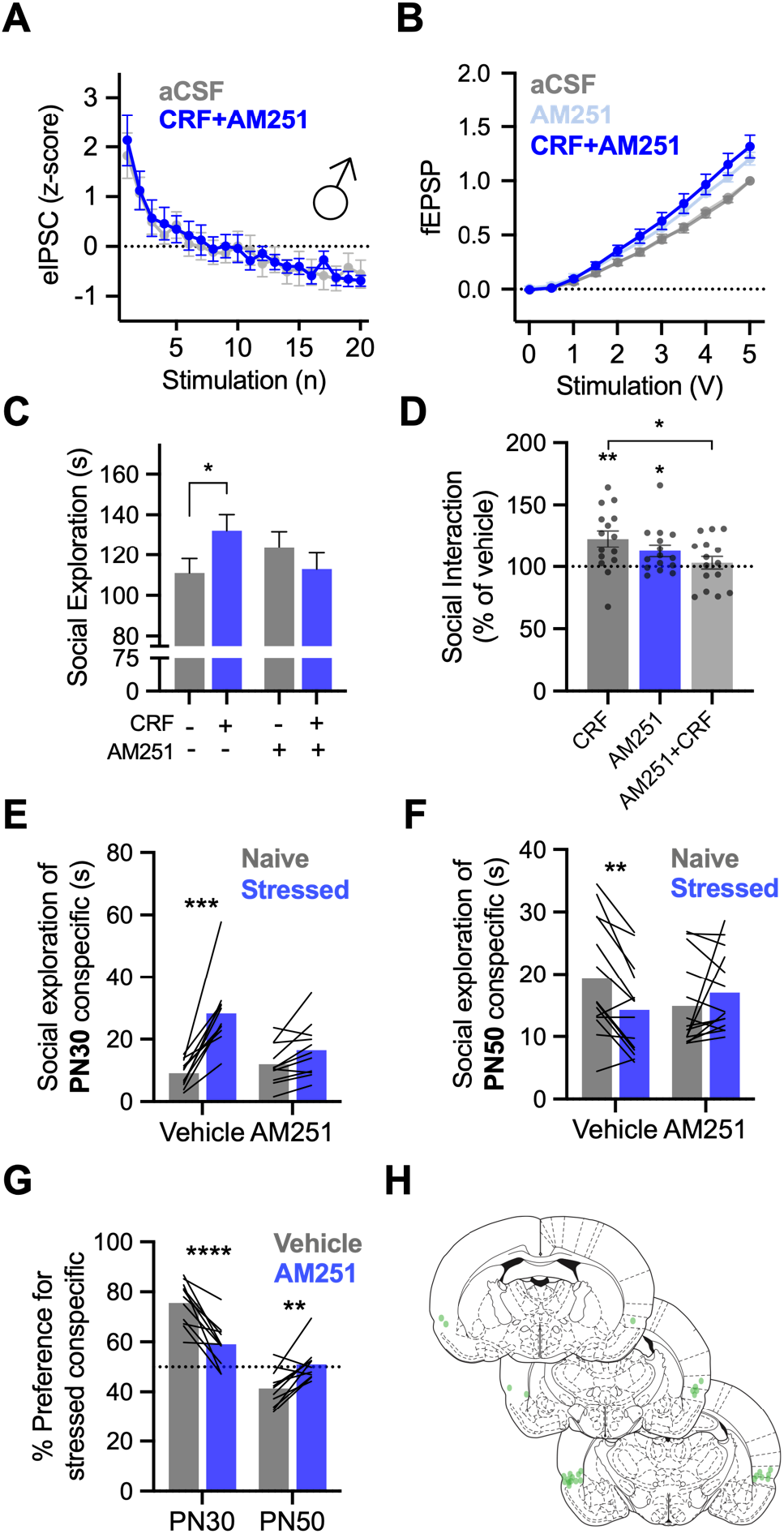
Cannabinoid 1 receptor is necessary for the behavioral and synaptic effects of insular CRF. **A**. fEPSPs recorded from insular cortex slices (n=8) were insensitive to 300 nM CRF when applied with CB_1_ receptor antagonist AM251 (2 *μ*M), F_Voltage × Drug_ (1,11) = 2.992, P = 0.0959. **B**. In voltage clamp recordings of insular cortex pyramidal neurons (n = 9), AM251 prevented the inhibition of eIPSCs previously caused by CRF (50nM) application, F_Drug_(1, 8) = 0.0157, P = 0.9032. **C**. In a 5 minute social interaction test with male rats (n = 15) CRF injected to the insular cortex increased exploration of juvenile conspecifics (P = 0.0270). CRF given in combination with AM251 did not increase social exploration, F_CRF × AM251_ (1, 14) = 9.102, P = 0.0092. **D**. For comparison, raw social interaction times from Panel C are shown as percent of time relative to the no drug condition. CRF increased interaction (one-sample t-test compared to 100%, t(14) = 3.422, P = 0.0041, AM251 alone increased interaction, t(14) = 2.620, P = 0.0202, but CRF given with AM251 did not differ from vehicle levels, t(14) = 0.5935, P = 0.5623. Importantly, social interaction was greater with CRF alone than in combination with AM521, (P = 0.0110, Sidak test after significant one-way ANOVA, F(2, 28) = 3.705, P 0.0374). **E**. In SAP tests with juveniles (n = 11), AM251 prevented the formation of a preference for stressed juvenile conspecifics, F_Drug × Stress_(1, 9) = 22.53, P = 0.0010, such that the preference for stressed juveniles present during vehicle testing (P < 0.0001) was eliminated during AM251 testing (P = 0.0940). **F**. In SAP tests with adult conspecifics (n = 16), AM251 eliminated the preference of test rats for naive adult F_Drug × Stress_(1, 13) = 19.93, P = 0.0006. A significant preference for naive adults was present in vehicle test rats (P = 0.0013) which was not present in AM251 rats (P = 0.1691). **G**. Percent preference for stressed conspecifics was significantly altered by a combination of AM251 treatment and age of conspecific F_Age × Drug_(1, 20) = 43.12, P < 0.0001. Specifically, the percent preference for stressed juveniles was significantly reduced (P < 0.0001) while the percent preference for stressed adults was significantly increased (P = 0.0054). **H**. Cannula placements of all animals in the experiments contained in Fig. 4. Bar graphs indicate mean with individual replicates, line graphs mean (+/− SEM). *P < 0.05, ** P < 0.01, *** P < 0.001, **** P < 0.0001 (Sidak’s tests).

### Sex differences exist in CRF_1_ and CB_1_ mRNA expression and cellular distribution in the insular cortex

The pattern of sex-specific effects of CRF on physiology and social behavior led us to hypothesize that sex differences exist in the relative amount and cellular distributions of the CRF_1_ and CB_1_. Using qPCR on insular cortex micro punches we found greater CRF_1_ mRNA in female rats (n=6) compared to males (n=8), t(12)=2.728, p=0.018 and no difference in CB_1_ mRNA expression, t(14)=0.090, p=0.929, Fig. 5A). We also attempted to quantify CRF (*crh*) and CRF type 2 receptor (*crhr2*) mRNA in these samples, but they were not detectable, which is consistent with prior reports ^28,29^. That females expressed more CRF_1_ mRNA than males is inconsistent with the electrophysiology and behavioral studies in which CRF had little or no effect in females. To understand the functional sex differences we turned to RNAScope fluorescent *in situ* hybridization, to visualize and colocalize mRNAs of glutamatergic neurons (*vGlut1*), CRF_1_ (*crhr1*) and CB_1_ (*cnr1*) in 20*μ*m sections of insula from 8 male (Fig. 5D) and 8 female (Fig. 5E) adult rats. The number of mRNA positive cells were quantified from tiled image stacks of the left and right posterior insula. Males had more DAPI nuclei labeled with CRF_1_ and CB_1_ mRNA (Fig. 5B). Looking at putative glutamate neurons (vglut1 positive cells, Fig. 5C), there was a trend for greater CRF_1_ in males, but this did not reach significance (p=0.067). However, in males we found more glutamate neurons co-localized with CB_1_ and more glutamate neurons co-localized with both CRF_1_ and CB_1_ mRNAs than females.

**Fig. 5.**
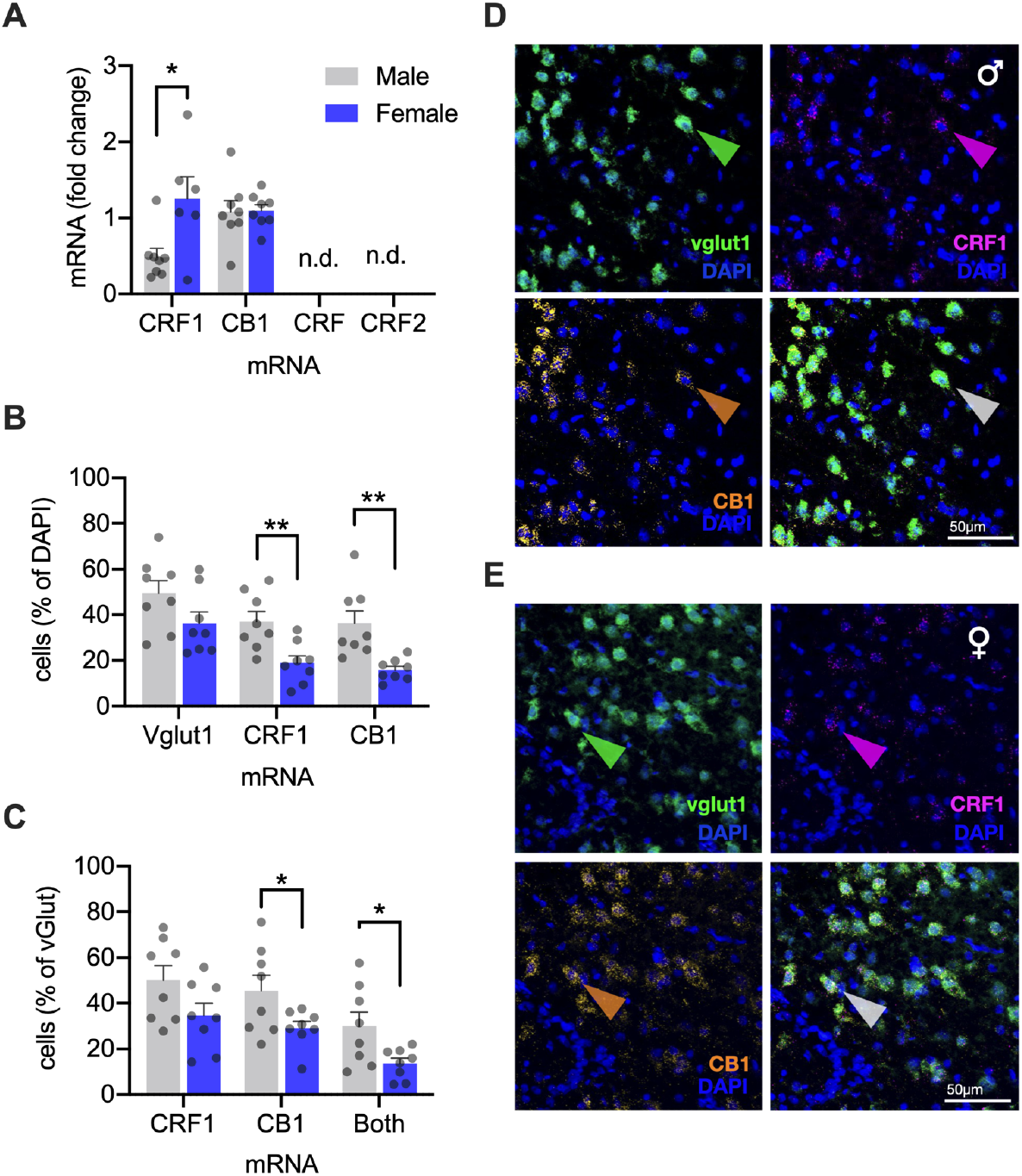
Cellular distribution of CRF_1_ and CB_1_ receptor mRNA in the insular cortex. **A**. qPCR analysis of relative mRNA expression revealed greater CRF_1_ mRNA in females compared to males t(12)=2.728, P = 0.183, and equal CB_1_ mRNA across sexes t(14) = 0.090, P = 0.929. **B**. RNAScope was performed for CRF_1_, CB_1_, and vglut1 mRNAs. Fluorescent grains were counted in the left and right hemispheres of the posterior insular cortex. The total number of cells was determined by counting DAPI nuclei in each hemisphere. Nuclei containing 3 or more fluorescent grains were considered mRNA expressing cells and shown as the % of the total cells. The number of vglut1 cells was equal between sexes, but the portion of cells expressing CRF_1_ F_Sex_(1, 14) = 11.19, P = 0.005, or CB_1_, F_Sex_(1, 14) = 13.07, P = 0.003, mRNA was approximately double in males compared to females. **C**. Looking at expression of CRF_1_ and CB_1_ mRNA in vglut1 cells, shown as a percent of the total vglut1 cells per hemisphere, male rat sections contained more CB_1_ mRNA expressing vglut cells, and more vglut cells expressing both CB_1_ and CRF_1_ mRNA, F_Sex_(1, 14) = 4.489, P = 0.044). The number of vglut cells expressing CRF1 mRNA was greater on average in males than females, but did not reach significance F_Sex_(1, 14) = 3.944, P = 0.067. In males, there were more vglut cells co-localized with both CRF_1_ and CB_1_, F_Sex_(1, 14) = 6.576, P = 0.023. **D-E**. Representative digital photomicrographs of RNAScope *in situ* hybridization and fluorescent visualization of DAPI and vesicular glutamate transporter 1 (*vglut1)*, CRF_1_ (crhr1), and CB_1_ (cn1r) mRNA from male (A) and female (rats) insular cortex coronal sections (20um, n=8/sex). Sections were selected from subjects nearest to the mean values for CRF_1_ + DAPI colabeling. Colored arrowheads indicate cells with coexpression of all three mRNAs with DAPI. Bar graphs indicate mean (+SEM) with individual replicates. *P < 0.05, ** P < 0.01, *** P < 0.001, **** P < 0.0001 (Sidak’s tests).

## DISCUSSION

We sought to identify neural mechanisms by which social stress signals influence social decision making. We focused on the hypothalamic neuropeptide CRF, which is released during social interactions with stressed conspecifics ^10^, and the posterior insular cortex (IC), a structure needed for social affective decision making ^22^ and a novel site of action for CRF. A combination of electrophysiology, pharmacology, behavior and molecular experiments revealed a sex-specific role for CRF as a modulator of insular synaptic physiology and social behavior. In males only, CRF caused a reduction in presynaptic inhibitory tone, likely via release of retrograde endocannabinoids acting at presynaptic CB_1_ receptors. Behaviorally, insular CRF injections increased social interaction and both CRF_1_ and CB_1_ receptor antagonists impaired social decision making in a social affective preference test. To understand the basis for the sex difference we used qPCR and *in situ* hybridization to describe the relative expression and cellular distribution of CRF_1_ and CB_1_ mRNAs. Although we found more overall CRF_1_ mRNA in females, we found that males had more CRF_1_ positive cells and increased CRF_1_ and CB_1_ colocalization on glutamate neurons. In sum, these results add to our understanding of CRF as a neuromodulator, integrate CRF into a social-decision making process, and have important sex-specific implications for understanding the neurobiology of social cognition and psychopathology.

The depolarizing effect of CRF on membrane potential is consistent with the G_S_-protein signaling and cAMP modulation of cation channels ^30^. However, the dampening effects of CRF on the AP, reduction of ADP and the increase in rectification ratio suggest additional effects of CRF via modulation of ion channels. Specifically, CRF can reduce T-type voltage-gated calcium channels ^31^ and modulate voltage-gated potassium channels ^32^. These intrinsic effects of CRF would predict a reduction in firing rate which was evident when comparing the spike rate of a 250pA current injection (Table 1) and also found in hippocampal recordings ^32^. These results are not likely the result of diffusion or cell dialysis effects during recordings because under identical circumstances and recording times, these parameters are stable ^22^. The most robust effect of CRF was depolarization of the membrane potential which we predicted would lead to augmentation of excitatory synaptic transmission. When looking at synaptic measures, however, sex-specific effects of CRF become clear. In males only, CRF augmented fEPSPs in a dose- and CRF_1_-dependent fashion. The synaptic effect of CRF also depended upon GABA_A_ receptors, suggesting a more complex mechanism of action for CRF than direct modulation of principal neurons via CRF_1_. We further investigated this in evoked IPSCs as a more direct measure of GABAergic tone and found a CRF_1_ dependent reduction in eIPSC amplitude in males, but not females. Because depolarization alone is sufficient to inhibit presynaptic GABAergic transmission via retrograde endocannabinoids and the CB_1_ receptor ^21^ we tested the dependence of CRF effects on CB_1_ and found that both the augmentation of fEPSPs and suppression of eIPSCs were prevented by pretreatment with a CB_1_ receptor inverse agonist. Interestingly, the forgoing effects of CRF on excitatory and inhibitory insular synaptic transmission were only evident in male rats. These results suggest that CRF action within the insular cortex begins via CRF_1_ mediated depolarization of pyramidal neurons which then release endocannabinoids which, in turn, cause suppression of presynaptic GABAergic neurons accounting for increase in excitatory transmission in male insula. Although glucocorticoid receptor stimulation is known to modify inhibitory tone via endocannabinoids ^33^, to our knowledge this is the first evidence of the stress peptide CRF exhibiting CB_1_-dependent effects on presynaptic inhibition. An important goal of future work is to understand the molecular cascade mediating this effect, the type of presynaptic cells affected by endocanabinoids and the sex differences in these systems that result in male-specific effects.

Detecting socioemotional cues and using this information to inform whether to approach or avoid others is a vital aspect of social behavior. The mechanism underlying how this social information is gathered and used has become a recent focus with previous work showing important contributions of the PVN generated CRF and the insular cortex. The discovery by Sterley et al., (2018) that exposure to stressed conspecifics potentiated PVN CRF neurons inspired the current study. CRF receptors are distributed across many structures critical to social behavior ^26^ and the observer stress response, a form of emotion contagion, may be elementary to more complex social cognition that involves a distributed social decision-making network ^34^. Here we show that CRF can contribute to social behavior as insular injections increased social investigation of both juvenile and adult conspecifics, but only in male test rats. In the social affective preference (SAP) test in which experimental rats may choose to interact with either stressed or naive conspecifics, we typically find a preference for interaction with stressed juveniles but avoidance of stressed adults ^22,22,35^. Here, the CRF_1_ antagonist prevented this pattern suggesting that an activation of the CRF acute stress response, as reported by Sterely et al., likely occurs in the experimental rat during the SAP test. Our results are consistent with a finding in mice that exposure to synthetic fox odor, a potent social stressor both potentiates prefrontal cortex excitatory synaptic transmission and increases defensive behaviors in a CRF_1_ dependent fashion ^36^. Further, the mechanism of CRF action *in vivo* may be similar to what we first report in the acute slice preparation because both the behavioral effects of CRF on social interaction and SAP behavior were prevented by the CB_1_ inverse agonist. The results raise several interesting new questions. Specifically, what is the role of CRF in social cognition? One possibility is that CRF contributes to negative affect encoding or emotional states such as fear. To test this we injected CRF to the insula during fear conditioning, or prior to recall using a procedure that results in an insula-dependent conditioned freezing response ^37^, but CRF had no effect (Sup. Fig. 3). Alternatively, CRF may augment information throughput among the social decision making, or salience networks during acute stress ^20^. Thus, interference with CRF in the insula, a highly connected node within this network, may be sufficient to disrupt the coordination of information processing during social interaction. As such, it would appear that CRF in the insula, and perhaps other regions, is a necessary part of the neurochemical milieu for social stress detection which shapes current, and potentially future, social decision making.

The CRF system is immensely complex, comprising of many sex, brain region and cell type specific differences and functional contributions to behavior. As such, while the finding that CRF in the insula acts in a sex-specific manner to augment social behavior and drive social decision making in males is novel, the fact that a sex difference exists is not surprising. Here we began to understand the molecular basis for the sex difference by quantifying the relative expression and cellular distribution of CRF_1_ and CB_1_ mRNAs. Interestingly, qPCR revealed a greater relative amount of insular CRF_1_ mRNA in females compared to males and no difference in relative CB_1_ expression between males and females, indicating that the synaptic and behavioral sex differences found in this paper are not due simply to less available CRF_1_. These results contrast some prior reports which find greater CRF_1_ mRNA in prefrontal cortex of males ^38^ but it is difficult to generalize across regions as CRF_1_ receptor binding varies in sex specific ways with female adult rats tending to have more binding than males ^39^. More CRF_1_ in females is directly at odds with the present observations and suggests that there are further sex differences in either the distribution of CRF_1_ or CB_1_ receptors across cortical cell types, or the intracellular signalling cascades driven by CRF binding in males that are distinct from females ^40^. While the latter requires directed research, here we found more CRF_1_ and CB_1_ mRNA containing nuclei and glutamatergic neurons in males compared to females. These results illuminate a mechanism for sex specific effects. First, more glutamatergic neurons in males are sensitive to CRF providing a larger population of neurons to be depolarized and affect GABAergic tone. Second, CB_1_ is far more prevalent in males, which confirms other reports ^41^ which may account for the attenuation of presynaptic eIPCSs and dependence of CRF effects on CB_1_ in males. While these observations give some clarity to the organization of this system in the male insula, it remains possible that the distribution of these receptors varies in sex and cell-specific ways within the cortical interneuron subtypes which is an exciting and clinically relevant direction for future research.

An explosion of fMRI studies in recent years implicates the insula in an impressive range of cognitive processes ^42^, including emotion recognition ^6^, salience detection ^2^, pain ^7^, drug craving ^5^ and so on ^4,43^. In social disorders like autism spectrum disorder and schizophrenia, abnormal insular activity is a reliable correlate of social faculties like emotion detection and empathy ^18^. In studies focusing on insular cortex in PTSD, insula activity is associated with anticipatory fear and anxiety ^44–46^ and acute stress increases insula activity and functional connectivity to the salience network ^47,48^. Although the current study investigated CRF in the insula in an acute time frame, stressor exposure often occurs many days, months or years before stressor induced symptoms emerge. Perhaps the most exciting clinical implication of the current study is that we provide a mechanism by which acute stressors could augment plasticity or attention processes within the insular cortex that may cascade into abnormalities in fear, anxiety, and social functions. Clinically significant future research must investigate how severe and chronic stressors affect insula and consider CRF and endocannabinoids as therapeutic candidates for early intervention.

This study describes a mechanism by which CRF acts in the insula to increase excitability leading to changes in social decision making. Electrophysiology and pharmacology suggests CRF acts at CRF_1_ on principle neurons, which then release endocannabinoids in turn acting at CB_1_ on presynaptic neurons. This leads to a presynaptic inhibition of GABA interneurons leading to an increase in excitatory synaptic efficacy which drives social behavior, affect discrimination and social decision making. The insula is important to many behaviors that are shaped by stress. As such, CRF in the insula may play an important role in the development and pathobiology of a wide range of psychiatric diseases. Indeed, traumatic stress is now being linked to later enhancement of processing within the intrinsic salience network ^9,49^ and so is a new focus for intervention in PTSD ^50^ and other stress related psychoses.

## METHODS

### Animals

Male and Female Sprague-Dawley rats were obtained from Charles River Laboratories (Wilmington, MA) at either age PN45 (test rats and adult conspecifics) or PN21 (juvenile conspecifics). All animals were allowed to acclimate to the vivarium for a minimum of seven days prior to any experimental or surgical procedures. Rats were housed with two or three same-sex conspecifics and given access to food and water *ad libitum*. The vivarium was maintained on a 12 h light/dark cycle and all behavioral tests occurred within the first four hours of the light cycle. All procedures and animal care were conducted in accordance with the NIH *Guide for the care and use of laboratory animals* and all experimental procedures were approved by the Boston College Institutional Animal Care and Use Committee.

### Electrophysiology

#### Solutions

Standard solutions were used for artificial cerebrospinal fluid (aCSF) and recording solutions ^51^ and all reagents were purchased from ThermoFisher, Sigma or Tocris. aCSF recording composition was (in mM) NaCl 125, KCl 2.5, NaHCO_3_ 25, NaH_2_PO_4_ 1.25, MgCl_2_ 1, CaCl_2_ 2 and Glucose 10; pH = 7.40; 310 mOsm; aCSF cutting solution was: Sucrose 75, NaCl 87, KCl 2.5, NaHCO_3_ 25, NaH_2_PO_4_ 1.25, MgCl_2_ 7, CaCl_2_ 0.5, Glucose 25 and Kynureinic acid 1; pH=7.40, 312 mOsm. The internal recording solution consisted of (in mM) K^+^-Gluconate: 115, KCl 20, HEPES 10, Mg-ATP 2, Na-GTP 0.3, and Na-Phosphocreatine 10. pH = 7.30; 278 mOsm with 0.1% biocytin. Kynurenic acid 1 mM and SR-95531 2 µM were added to the recording aCSF to block synaptic transmission for intrinsic recordings. Synaptic blockers were omitted for fEPSP recordings, except where noted, and SR-95531 was excluded for evoked inhibitory postsynaptic current (eIPSC) recordings.

#### Insular Cortex Slices

Adult male and female rats were anesthetized by isoflurane, intracardially perfused with chilled (4°C) aCSF cutting solution and rapidly decapitated. The brain was sliced on a vibratome (VT-1000S, Leica Microsystems, Nussloch, Germany) in 300 µm coronal segments containing the insula. Slices were then transferred to oxygenated (95% O_2_, 5% CO_2_) aCSF cutting solution at 37°C for 30 minutes followed by 30 minutes at room temperature prior to any electrophysiological recordings.

#### Whole Cell Recordings

Active and passive membrane properties of deep layer insular cortex pyramidal neurons were determined in acute whole cell current clamp recordings. Experiments were conducted in pClamp 10 using an Axon 700B amplifier, headstage and DigiData1550 DAQ. Experimental protocols were exactly as described previously in Rogers-Carter et al. 2018. Patch-clamp electrodes were pulled (P-1000, Sutter Instruments, CA) from 1.5 mm outer diameter borosilicate glass (Sutter Instruments, CA) and filled with an intracellular solution. Electrode resistance was 3–5 MΩ in the bath and recordings were only included if the series resistance remained less than 30 MΩ with less than 10% change from baseline throughout the experiment. Slices were visualized using a 40x (0.75 NA) water immersion objective under infrared differential interference contrast imaging on an upright microscope (AxioExaminer D1, Zeiss, Germany). All recordings were obtained with an Axon 700B amplifier, DigiData1550, and pClamp 10 (Molecular Devices), using appropriate bridge balance and electrode-capacitance compensation. After achieving a whole-cell configuration, baseline recordings were made in aCSF until 10 minutes of stable baseline were observed, at which point 50nM CRF (human/rat, Cat. No 1151, Tocris) was added to the bath. The dose of 50 nM was selected after a pilot study using a range of doses from 50nM to 300nM, representative of the low ^52^ and high ^36^ to doses found in the literature. While dose responses were seen at the level of the synapse (Fig. 2) no dose responses were seen in intrinsic measures between 50 nM and 300 nM concentrations and thus all patch-clamp experiments were completed at 50 nM. Analyses were performed using custom software written for Igor Pro (Wavemetrics Inc., Lake Oswego, OR). Active properties were quantified from single spikes by holding the neuron at −67 mV, and 2.5 ms current pulses were injected to elicit a single AP. Passive properties were measured by holding the membrane potential at −67 mV and injecting 1 s current pulses through the patch electrode. The amplitudes of the current injections were between −300 pA and +400 pA in 50 pA steps. All traces in which APs were elicited were used to generate input-output curves as the total number of APs per second plotted against the injected current. A complete list of parameters is provided in Table 1. The burst ratio was computed as previously described as “burst current” ^53^ in which a larger ratio indicates more current required of a transient depolarization applied during the ADP to evoke an AP.

#### Evoked Field Excitatory Postsynaptic Potentials

Evoked field excitatory postsynaptic potentials (fEPSP) were recorded on a 6 × 10 perforated multielectrode array (MCSMEA-S4-GR, Multichannel Systems) and integrated acquisition hardware (MCSUSB60) and analyzed with MC_RACK software (version 3.9). Slices were placed onto the array and affixed by downward suction of the substrate through the perforated array by perfusion. CRF (50 or 300 nM), CRF_1_ antagonist CP-154526 (10 µM) and GABA_A_ inhibitor SR-95531 (2 µM) were dissolved in water or DMSO and then diluted to their final concentration in aCSF and bath applied to slices from above at 37°C.

A stimulating electrode was selected in the insular cortex and fEPSPs were recorded throughout the rest of the insular cortex following electrical stimulation before (baseline), during (drug) and after (wash) drug application. Stimulations ranged from 0-5 V and occurred in biphasic (220 µs) 500 mV increments. Each step in the I/O curve was repeated three times for a total of 33 stimulations in each of the three conditions. fEPSPs that showed clear synaptic responses and occurred close to the stimulating electrode (Fig. 2A) were normalized to the maximum 5 V response of the slice at baseline and channels from the same slice were averaged for group analysis across subjects. Slices received only one drug treatment.

#### Evoked IPSCs

To determine the effect of CRF on GABA_A_-mediated IPSCs, electrical stimuli were applied via a bipolar stimulating electrode; AMPA & NMDA receptors were blocked with 10 μm CNQX and 50 μm APV; action potentials were blocked intracellularly with the sodium channel blocker QX-314 (10mM) in the internal solution. Trains of 20 stimuli were applied at 5Hz to evoke inward eIPSCs were recorded from holding potential of −90mV as previously reported ^54^. eIPSCs were quantified as the peak amplitude observed in the 1ms post stimulation. Due to inter-recording variability in absolute current size, each recording was converted to a z-score using the mean and standard deviation of the baseline (aCSF) eIPSC peaks for analysis with ANOVA. Charge (area of IPSC and IPSC slope were also computed, normalized and analyzed the same way, the effect of CRF was the same with each of these approaches (data not shown).

### Surgical implantation of insula cannula and microinjection

While under isoflurane anesthesia (2-5% v/v in O_2_), rats were surgically implanted with bilateral guide cannula (26-gauge, Plastics One, Roanoke VA) in the insular cortex (from bregma: AP: −1.8 mm, M/L: ± 6.5 mm, D/V: −6.8 mm from skull surface) that were affixed with stainless steel screws and acrylic dental cement. Immediately following surgery, rats were injected with analgesic meloxicam (1 mg/kg, Eloxiject, Henry Schein), antibiotic penicillin (12,000 units, Combi-pen 48, Covetrus) and Ringer’s solution (10 mL, Covetrus). Rats were then returned to their homecage and allowed 7-14 days for recovery prior to behavioral testing.

CRF was first dissolved in DI water and diluted to 300 nM concentration in a vehicle of 0.9% saline. CRF antagonist CP-154526 (Sigma) was first dissolved in 100% DMSO before being diluted to 10 µM in a vehicle of 10% DMSO and 0.9% saline. AM251 (Tocris) was first dissolved in 100% DMSO prior to being diluted to 2 µM in a vehicle consisting of 10% DMSO and 0.9% saline. Injections were 0.5 µL/side for all drugs and were infused at a rate of 1.0 µL/minute with an additional one minute diffusion time after injection. After behavioral testing concluded rats were overdosed with tribromoethanol (Sigma) and brains were immediately dissected and fresh frozen on dry ice. Brains were sectioned on a cryostat at 40 µm, slices were then mounted onto gel-subbed slides (Fisher) and stained with cresyl-violet to verify cannula placements by comparing microinjector tip location to the rat whole brain stereotaxic atlas. Rats were excluded from all analyses if their cannula placements were found to be outside the insula or if both of their cannulas were occluded prior to injection.

### Social Exploration

One on one social interaction tests (SE) were completed in a quiet room under white light. Each test rat was assigned to a separate plastic cage (18 cm × 24cm × 18 cm) with shaved wooden bedding and a wire lid that was used throughout testing (between 2 and 4 social interactions depending on the experiment). Rats were moved from their housing cage to the testing cage prior to experimenting and allowed 1 h to acclimate to the testing cage. 30-40 minutes prior to testing, rats were microinjected with either drug or vehicle as described above. The test began with the introduction of either a juvenile (28 ± 2 days old) or adult (50 ± 2 days old) same-sex conspecific. Rats were then allowed to interact for five minutes and interactions were scored for social behaviors (sniffing, pinning and allogrooming) initiated by the test rat by an observer blind to treatment. Rats were tested in a within subjects design where they were tested on consecutive days. For CRF testing (Fig. 3A), rats received either vehicle or CRF in a counterbalanced order. For CRF1 antagonist, rats received four treatments (vehicle, CRF, CRF_1_ antagonist, or CRF + CRF_1_ antagonist) across four consecutive testing days counterbalanced in a latin square design. For AM251, rats received four treatments (vehicle, CRF, AM251, CRF + AM251) across four consecutive testing days using a latin square design for counterbalancing. Conspecifics were used more than once but no test rat was paired with the same conspecific twice.

### Social Affective Preference (SAP) test

The SAP tests allows for the quantification of social interactions initiated by a test rat towards either a stressed or unstressed conspecific, providing insight into the test animals discrimination of socioemotional affective cues. The SAP test occurs over four days (Fig. 3G) with days 1 and 2 serving as habituation days and days 3 and 4 serving as testing days. On day one the rat is placed into an empty clear plastic cage (50 cm × 40 cm × 20 cm) for one hour. On day two rats are placed into the cage for one hour prior to testing with two naive juvenile or adult conspecifics. Conspecifics are placed into opposite ends of the testing cage in a clear plastic 18 cm × 21 cm × 10 cm cage with one side made up of horizontal bars spaced 1 cm apart to facilitate interactions. The test rat is then allowed to interact with the conspecifics for five minutes and is scored for time spent nose-nose or nose-body sniffing and reaching for the conspecific through the bars. On days three and four 40 minutes prior to testing rats were microinjected with vehicle or CRF_1_ antagonist as described above in a counterbalanced, within subjects design. Also on days three and four, one of the conspecifics placed into the cage is stressed via two 5-second 1 mA footshocks in 60 seconds immediately before testing. Conspecifics were then placed in the chamber and scoring was done as described above by an observer blind to treatment. Interactions were also video recorded for scoring by a second observer to determine interrater reliability.

### Insular mRNA Quantification

To determine the relative expression of CRF_1_ (crhr1), CB_1_ (Cnr1) CRF (crh), and CRF_2_ (crhr2) mRNA we performed quantitative Taq-man reverse transcriptase polymerase chain reaction (qPCR) analysis on 1mm dia, ∼500μm thick insular cortex punches. Punches were collected on a freezing cryostat by slicing fresh frozen brains to a coronal plane equal to Bregma −1.5mm and depressing a 1mm brain punch tool (Stoelting) through the insula such that the base of the punch intersected the rhinal fissure. The resulting punches contained the extent of insula targeted for electrophysiology, microninjection and RNAscope analysis. Punches were kept on dry ice until storage at −80°C. As previously described ^55^, total RNA was isolated using the RNAqueous-4PCR Total RNA Isolation Kit (Applied Biosystems, Foster City, CA, United States) and processed per the manufacturer’s instructions. cDNA was synthesized using the High Capacity cDNA Reverse Transcriptase Kit (Applied Biosystems, Foster City, CA, United States). cDNA concentration was quantified using Nanodrop 2000 Spectrophotometer (Thermo Fisher Scientific), and samples were diluted accordingly. Amplification was performed using TaqMan Gene Expression Assays using gene-specific primers purchased from Thermo Fisher Scientific for *Crhr1* (Rn.10499), *Cnr1* (Rn.89774),. Reactions were performed in triplicate on a StepOnePlus Real-Time PCR System (Applied Biosystems) using standard cycling conditions, as recommended by the manufacturer. 4μl of diluted cDNA was placed in a 20μl reaction plate containing 16μl of master mix and 1x dilution of each primer. Reactions were performed with an initial holding stage of 50°C for 2 min and 95°C for 10 min, followed by 40 subsequent cycles of 15s at 95°C and 1 min at 60°C. The relative standard curve analysis method was used, with the threshold cycle (CT) (number of cycles required to reach detection threshold) determined for each reaction and the 2^-ΔΔC(t)^ method used to determine gene expression relative to the house-keeping control gene, β-actin (Thermo Fisher Scientific, Rn.94978). Data are expressed as mean fold expression relative to females.

### RNAScope *in situ* Fluorescent Hybridization

Coronal sections containing the posterior insular cortex (Bregma −1.8) were collected on a freezing cryostat at 20μm thick and mounted to SuperFrost+ slides and stored at −80°C until processing. RNAScope was performed according to the vendor’s instructions (ACDBio). Briefly, tissue was thawed, fixed and treated with a RNAScope cocktail including probes for CRF_1_ (*crhr1*, catalog #318911), CB_1_ (cb1, catalog #412501), vesicular glutamate transporter 1 (*vglut1, catalog #317001*) and DAPI. Probes were amplified and visualized with AMP1, AMP2, AMP3, AMP4, and DAPI, coverslipped with aqueous antifade media (Prolong Gold) and imaged on a Zeiss AxioImager Z2 microscope with a digital CCD camera (ORCA 3, Hamamatsu) using an Apotome2, 20x objective (N.A. = 0.8) and fluorescent filter cubes for DAPI (365 nm excitation, Zeiss filter 49), GFP (470/40 nm excitation, Zeiss filter 38 HE), DSRed (545/25 nm excitation, Zeiss filter 43 HE) and Cy5 (640/30 nm excitation, Zeiss filter 50). All image acquisition parameters (exposure, camera gain, and display curves were consistent for all samples. A series of multiplex, tiled mosaic images consisting of 9 z-series images per channel were stitched, deconvolved and maximum projections were saved for analysis in ImageJ. All channels were converted to binary and DAPI cells and vglut1, CRF_1_ and CB_1_ grains were detected using the particle counter tool. Trained observers counted the number of DAPI nuclei that were colocalized (a minimum of 3 overlapping, or adjacent grains) with each of the targets. The total number of nuclei (DAPI) and glutamate cells (Vglut+DAPI), and cells colocalized with CRF_1_, CB_1_ or both were determined. Cell counts were normalized by the total number of DAPI or total number of DAPI+vglut cells to compare the distribution of mRNAs with 2 way ANOVAs with sex as a between groups factor and side as a within-subject factor. The mean of left and right hemisphere counts are shown.

### Statistics

All statistics were run using Prism (Graphpad, version 8.0.3). Sample sizes were determined based on previous work for electrophysiological tests and based on previous work and power analyses for behavioral tests. Animals were randomly assigned to experiments and within subjects designs were used throughout. To control for potential test-order effects in repeated measure experiments, subjects were randomly counterbalanced (for 2 treatment tests) or ordered in a latin square design (for 4 treatment tests). Rats were only excluded from tests if their cannulas were not in the insula or if their cannulas were occluded prior to testing. To compare differences between mean scores of social interaction and electrophysiological endpoints we used t-tests and analysis of variance (ANOVA). Individual replicate data are provided in the figures. Data were checked for normality and transformed where appropriate via log transformation, final sample sizes are indicated in the Figure Legends. In most experiments, there were within-subjects variables, which were treated as such in the analysis (paired samples t-test or repeated measures ANOVA). Main effects and interactions were deemed significant when p < 0.05 and all reported post hoc test *p* values are Tukey or Sidak-adjusted, to maintain an experiment-wise risk of type I errors at a = 0.05.

## Author Contributions

Conceptualization, N.S.R., J.A.V., H.C.B., J.P.C.; Methodology, N.S.R., J.A.V., A.N., L.G., J.P.C.; Investigation, N.S.R., J.A.V., A.N., L.G., A.D., J.P.C.; Writing – Original Draft, N.S.R. and J.P.C.; Writing -- Revision & Editing, N.S.R. and J.P.C.; Funding Acquisition, J.P.C.

## Acknowledgements

The authors wish to thank Dr. Bret Judson, director of the Boston College Imaging Core, for training and assistance with all microscopy, Nancy McGilloway and Todd Gaines, administrators of the Boston College Animal Care Facility, for outstanding animal husbandry, Rahul Alturi, Benjamin Tramonte and Emma Fritsch for assistance with RNAscope, and Bridget Brady and Shanon Lee for help with MEA electrophysiology and behavioral tests. Funding for this work was provided by the Boston College Undergraduate Research Fellowship, the Gianinno Family, and the National Institutes of Health Grants MH119422 & MH109545.

---

**Supplemental Fig. 1.**
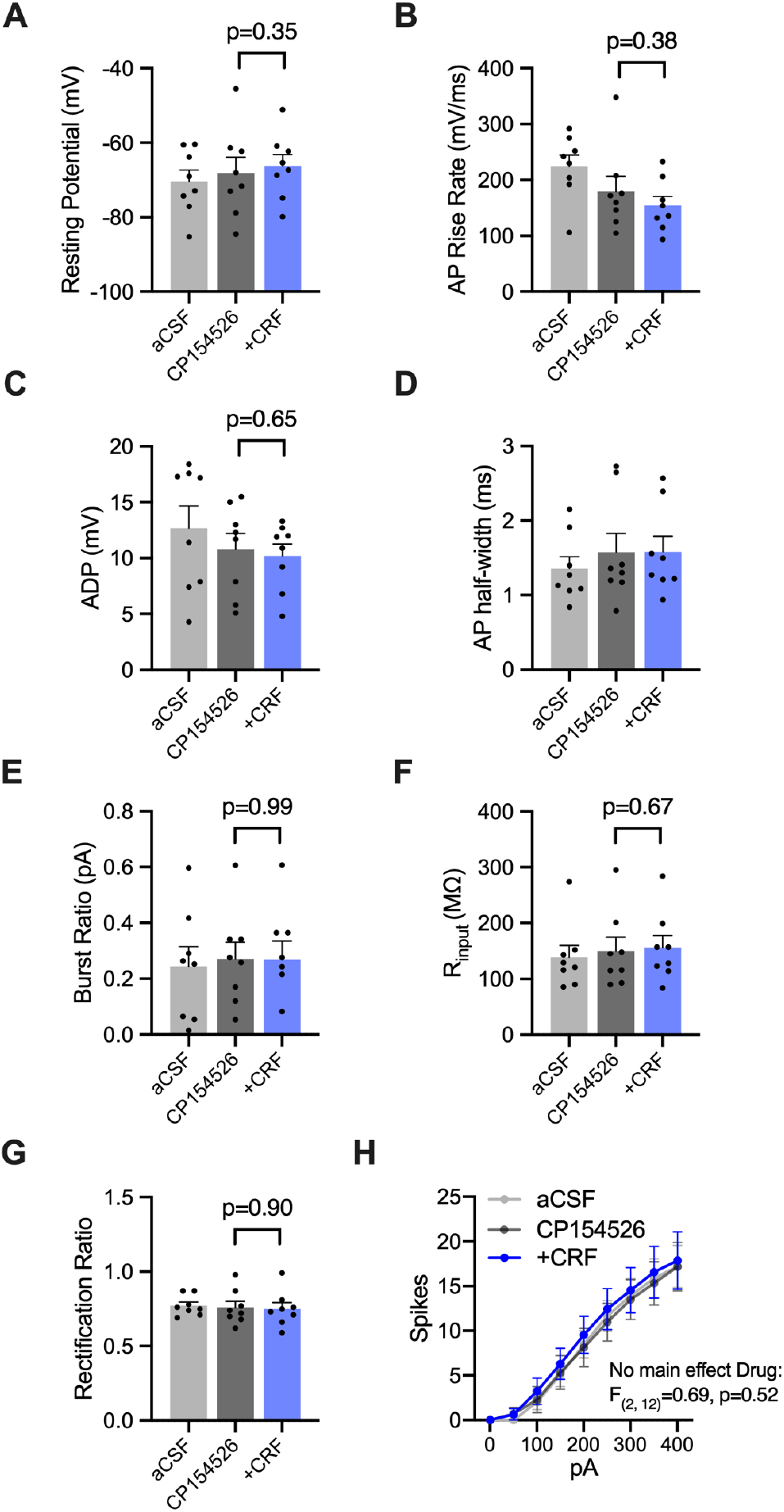
CRF does not alter intrinsic membrane properties in the presence of CRF_1_ receptor antagonist. Whole cell recordings were made in 8 male insular pyramidal neurons and intrinsic properties were determined exactly as in Figure 1 in regular aCSF, aCSF + CP154529 (10uM) and aCSD + CP154529 + CRF (50nM). The CRF_1_ antagonist prevented changes previously observed after CRF application (see Figure 1) including **A**. resting potential **B**. action potential rise rate **C**. after depolarization **D**. action potential half-width **E**. burst ratio **F**. input resistance, **G**. rectification ratio and **H**. firing rate. All measures were evaluated for drug effects with repeated measures ANOVA. There were no significant main effects of drug and no significant posthoc comparisons. P values are Sidak corrected.

**Supplemental Fig. 2.**
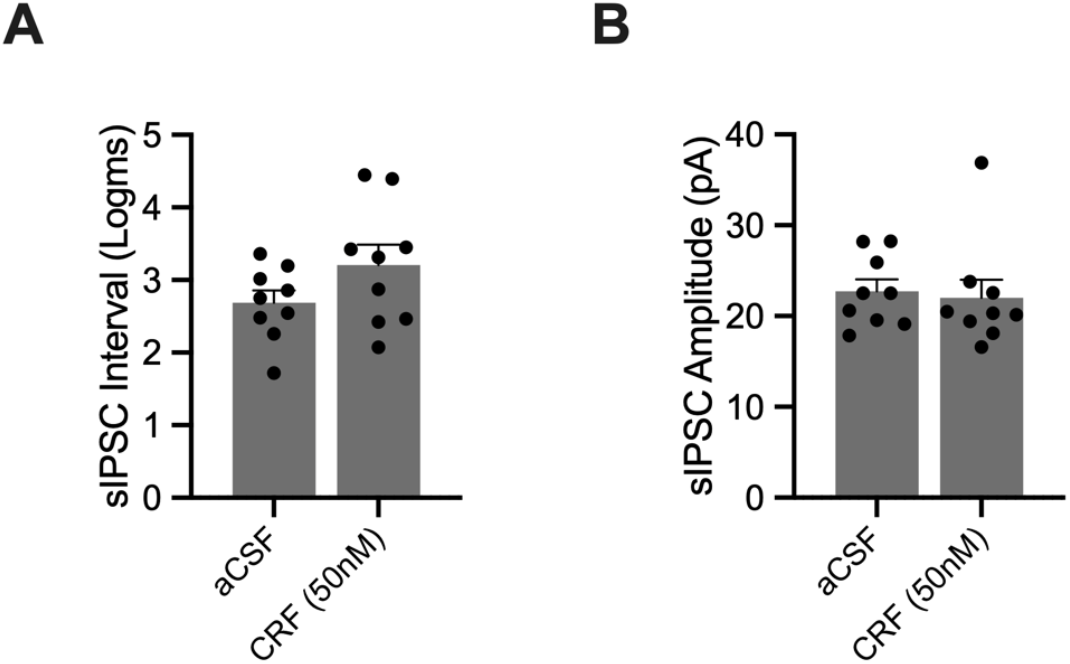
CRF did not alter spontaneous inhibitory postsynaptic currents (sIPSCs) in insular cortex pyramidal neurons. Whole cell voltage-clamp recordings were made and sIPSCs were recorded from 9 insula pyramidal neurons for 10 minutes in aCSF, and 10 minutes after application of CRF (50nM) using standard protocols (see Rogers-Carter, Varela et al., 2018). **A**. Spontaneous IPSC intervals were normalized with a log conversion and were not altered by CRF. **B**. Spontaneous IPSC amplitude was also not changed by CRF in the presence of CRF_1_ antagonist.

**Supplemental Figure 3.**
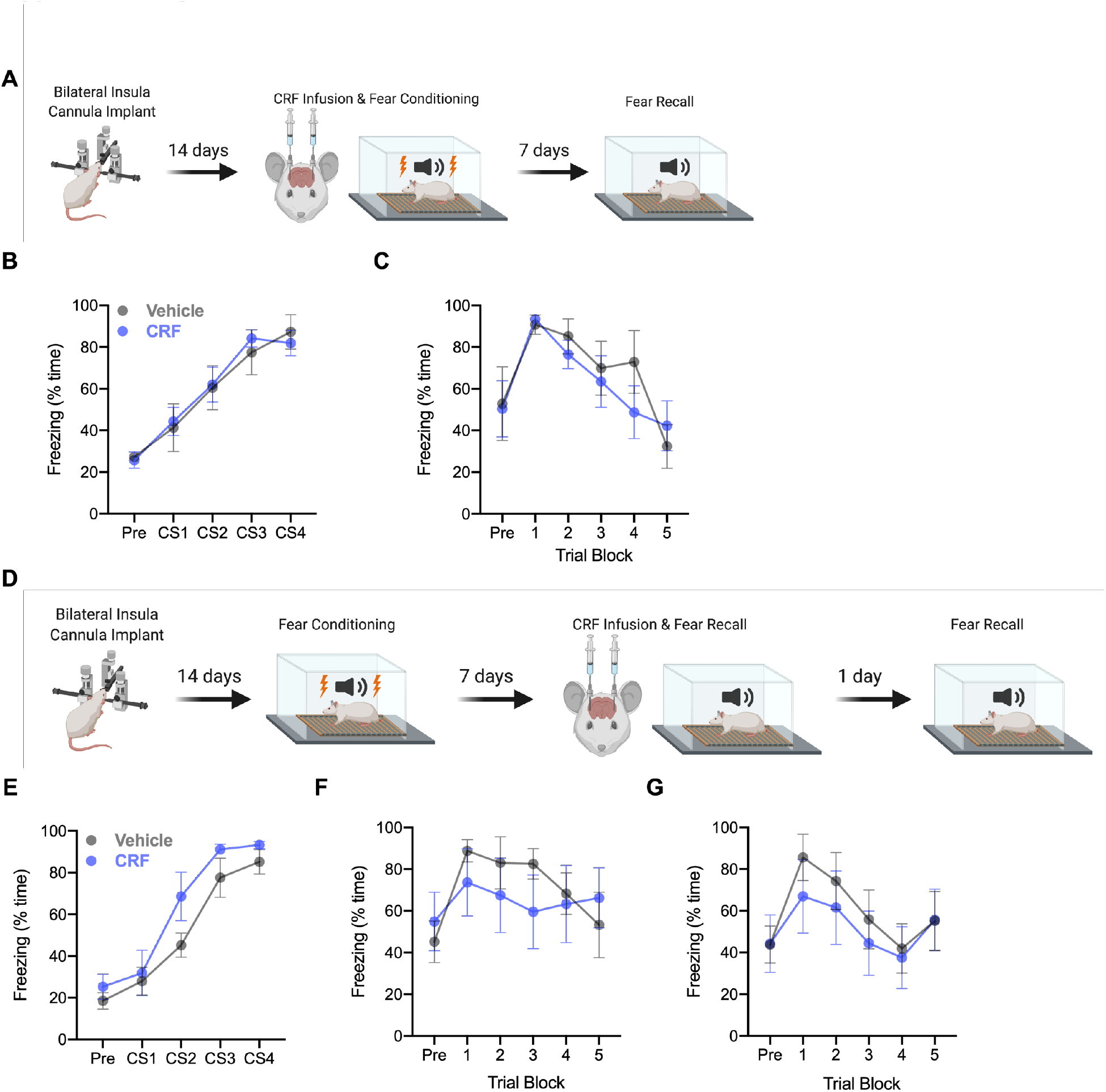
CRF in the insula does not affect fear learning or recall. **A**. Timeline of experiment for insular CRF on fear learning and recall. Male rats were bilaterally cannulated 14 days prior to undergoing fear learning by pairing a tone with a foot shock over 4 trials. 7 days after fear learning, fear recall was tested by measuring freezing in response to the shock-paired tone. **B**. Fear expression was not affected by microinjection of CRF (n = 6) or saline (n = 5) 40 minutes before conditioning. **C**. Later fear recall was not affected by CRF microinjection prior to fear learning. **D**. Timeline of experiment to determine if CRF microinjection prior to fear recall affected freezing. Male rats were implanted with bilateral insula cannula and underwent fear learning 14 days later as in A. 7 days following fear learning rats were microinjected with CRF and tested for fear recall. Rats were then retested for fear recall one day later. **E**. Vehicle (n = 6) and CRF (n = 6) rats showed no difference in freezing expression during conditioning as in B. **F**. Rats showed no change in freezing during fear recall following CRF microinjection. **G**. Rats showed no change based on CRF microinjection in a second trial of fear recall one day later. Methods and apparatus for fear conditioning were previously described in Foilb et al., 2016 *Neurobiology of Learning and Memory*.

